# Predicting fitness related traits using gene expression and machine learning

**DOI:** 10.1101/2024.02.14.580307

**Authors:** Georgia A. Henry, John R. Stinchcombe

**Affiliations:** Department of Ecology & Evolutionary Biology, University of Toronto, Toronto, Ontario, Canada; Koffler Scientific Reserve at Joker’s Hill, University of Toronto, King, Ontario, Canada; Swedish Collegium for Advanced Study, Uppsala University, Uppsala, Sweden

## Abstract

Evolution by natural selection occurs at its most basic through the change in frequencies of alleles; connecting those genomic targets to phenotypic selection is an important goal for evolutionary biology in the genomics era. The relative abundance of gene products expressed in a tissue can be considered a phenotype intermediate to the genes and genomic regulatory elements themselves, and more traditionally measured macroscopic phenotypic traits such as flowering time, size, or growth. The high-dimensionality, low sample size nature of transcriptomic sequence data is a double-edged sword, however, as it provides abundant information but makes traditional statistics difficult. Machine learning has many features which handle high-dimensional data well and is thus useful in genetic sequence applications. Here we examined the association of fitness-components with gene expression data in *Ipomoea hederacea* (Ivyleaf Morning Glory) grown under field conditions. We combine the results of two different machine learning approaches and find evidence that expression of photosynthesis-related genes is likely under selection. We also find that genes related to stress and light response were overall important in predicting fitness. With this study we demonstrate the utility of machine learning models for smaller samples, and their potential application for understanding natural selection.

## Introduction

Natural selection is a ubiquitous evolutionary force which acts on the phenotypes of individuals in a population, resulting in genetic changes if heritable variation exists for the traits under selection. Understanding the shape and strength of natural selection typically requires observers to choose specific traits to measure which may or may not be the subject of selection (Lande and Arnold 1983). The type of traits studied has been found to influence the strength and temporal constancy of selection, with fecundity traits experiencing more consistent and strong selection than survivorship (Hoekstra et al. 2001; Kingsolver et al. 2001). Trait correlations and pleiotropy can additionally make detecting selection statistically difficult due to their potential to reduce power (Hersch and Phillips 2004). With high-throughput phenotyping, such as metabolomics or gene expression profiling, investigators often obtain a vast number of highly correlated traits, making traditional phenotypic selection analysis difficult or impossible. Machine learning modelling and regularization techniques, which can alleviate issues of multicollinearity and overfitting due to an overabundance of explanatory variables, now make it possible for investigators to avoid having to decide which traits may be important to selection, and instead observe them from the data. Here, we use machine learning modelling to estimate natural selection on gene expression, using estimates of fitness and gene expression obtained from a field experiment with the Ivyleaf morning glory (*Ipomoea hederacea*).

Gene expression can be viewed as an intermediate between the genome and traditionally observable traits, and reflects both the underlying genetic makeup and the environment (Liao and Weng 2015; Josephs 2021). Quantifying gene expression through RNA sequencing of tissue yields a less biased sampling of phenotypic traits compared to an observer’s opinions on what might be an important trait (Rockman and Kruglyak 2006; Josephs 2021), although investigators must still decide on growth or rearing environments, as well as the tissues and developmental stages to be sampled. RNA-sequence data is also highly multivariate, in that it captures a sample of all the genes being expressed in a tissue at the moment of sampling, yielding a broad representation of many phenotypes simultaneously. Although gene expression was initially expected to largely be under stabilizing selection (e.g, from mutation accumulation studies, Rifkin et al. 2005; also see Gilad et al. 2006), recent results suggest that there may directional selection on genes’ expression, especially when individuals are farther from the fitness optima. For example, Groen et al. (2020) measured selection on gene expression under drought and standard field conditions in domesticated rice (*Oryza sativa*). Using univariate approaches, they were able to estimate selection differentials for individual genes’ expression, finding generally weak selection. Selection was stronger under drought conditions, with earlier flowering time, modulated apparently by a single gene, allowing plants to “escape” the effects of drought (Groen et al. 2020). They were additionally able to identify significant selection on genes related to photosynthesis and growth, using a multivariate approach where they regressed fitness on principal component (PC) scores that summarized expression of many genes at once. Several investigators have constructed gene co-expression modules (Palakurty et al. 2018, Josephs et al. 2020, Brown and Kelly 2021), and then related multivariate estimates of expression of the entire module to phenotypic traits that are either indicative of overall plant performance or size, or are traits likely to be under natural selection such as components of life history. These experiments have found the expression of multiple genes in a module to be significantly associated with traits likely to be under natural selection in populations, thus implicating the expression levels of genes in the modules as potential targets of natural selection. Collectively, these studies illustrate how measuring selection on gene expression may facilitate identifying genes or pathways which are important for fitness (Price et al. 2022).

The dominant method of characterizing contemporary phenotypic selection is the Lande-Arnold (1983) regression approach, and variants of it (Kingsolver and Schemske 1991, Rausher 1992). In this approach, estimates of relative fitness are regressed on multiple, potentially correlated phenotypes. The resulting coefficients, termed selection gradients, measure the direct effect of the phenotype on relative fitness, accounting for the effects of other potentially correlated traits included in the model (Lande and Arnold 1983). While this approach has yielded thousands of estimates of natural selection (see Kingsolver et al. 2012), it faces fundamental limitations with gene expression data. Transcriptomic sequencing data is typically plagued by the “small *n*, large *p*” problem, where the number of “features” (estimates of expression) far exceeds the sample size. As a consequence, the multiple regression approach to measuring selection is untenable, as there are vastly more phenotypes than observations. Past studies seeking to measure contemporary selection on gene expression have thus used data reduction techniques such as PCA (Groen et al. 2020) or the construction of gene coexpression modules, whose overall expression is itself summarized with PCA (eigen-expression; Josephs et al. 2020, Brown and Kelly 2021). While these approaches allow estimates of selection on PC scores – of either the whole transcriptome or a co-expression module – they do not allow direct or indirect selection on an individual gene’s expression to be distinguished. Consequently, one of the dominant benefits of the Lande-Arnold (1983) approach is lost. Perhaps because of the “small *n*, large *p*” problem or the difficulty of interpreting selection on PC scores (Chong et al. 2018), we have dramatically fewer estimates of phenotypic selection on gene expression, especially compared to morphology, behavior, and life history.

Machine learning tools are often well-suited to handling the “curse of dimensionality” due to intrinsic and extrinsic regularization and are thus useful in the analysis of sequence data (Schrider and Kern 2018; James et al. 2021), and may be useful for measuring selection on multivariate measures of gene expression. Machine learning typically includes some type of loss function which serves to balance under- and over-fitting of the model, and as such observations with extreme values are not overly influential. The data is split into a training set (which is used to fit the model) and a testing set of data, which can then be used to evaluate the model performance, which can buffer against some overfitting by reducing the amount of statistical noise being fit to the model. Additionally, feature selection either through manual evaluation and pruning (as in Principal Component Regression), or through intrinsic model behaviour (as in LASSO regression or Decision Trees), can alleviate problems due to too many explanatory variables. Regularized estimates are biased (i.e., no longer unbiased, least squares estimates) but often lead to more accurate overall prediction (see Morrissey 2014, Sztepanacz and Houle 2024). Generally, machine learning tools focus on optimizing prediction rather than estimation of parameter values within a model (Schrider and Kern 2018; James et al. 2021; Greener et al. 2022) and thus are valuable for understanding complex nonlinear relationships between variables. While there has been some exploration of using regularized regression to estimate phenotypic selection (Morrissey 2014, Sztepanacz and Houle 2024), to date applications of machine learning to studies of phenotypic selection, especially for gene expression, have been limited.

We used gene expression data from 96 *Ipomoea hederacea* individuals grown under natural field conditions to detect important fitness-related genes using various machine learning methods. We examine the union of unsupervised and supervised approaches. Unsupervised machine learning models are those in which the observations are not “labeled” with a response variable, and supervised models in contrast use observations which are “labeled” with a response variable. We first use principal component analysis on gene expression as an unsupervised dimensionality reduction method followed by linear regression of relative fitness on principal component scores. We then trained a supervised classification model to predict whether an individual set seed based on our gene expression data. In combining disparate methods, we sought to find consistently important candidate genes or biological processes which are likely contributing to differences in fitness among individuals in the field. We investigated the predictive ability of these machine learning approaches and extracted the genes whose expression contributed most to the model fit (i.e. those associated most strongly with fitness differences). We then used Gene Ontology (GO) analyses to deepen our understanding of the types of genes which were significantly enriched in each model and the biological processes commonly found to be important between the models.

## Results

### Selection differentials for gene expression

We initially estimated selection differentials for gene expression, using two analyses. First, for each gene, we estimated the difference in expression between individuals that set seed (the after-selection cohort, *z\**) and the total population, regardless of seed set (the before selection cohort, *z*). We found a wide distribution of selection differentials centered on zero (Fig. 1A), suggesting that total selection to increase or decrease expression appears equally common, and that some genes are under strong selection to increase or decrease expression. While these analyses match the structure of the classification algorithms, we used for machine learning (*see below*), they ignore the differential fecundity of individuals that set seed. We secondly estimated selection differentials as the covariance between expression and relative fitness, which includes differential fecundity (Fig. 1B), finding much the same pattern as in the binary fitness selection differentials, although the distribution is slightly broader. The two estimates of selection differentials were similar (Fig. 1C), indicating that differential fecundity is a key component of selection on expression. Because selection differentials measure total selection on a phenotype and cannot distinguish direct selection or indirect selection via correlated traits, we next turned to selection gradients for gene expression.

**Figure 1:**
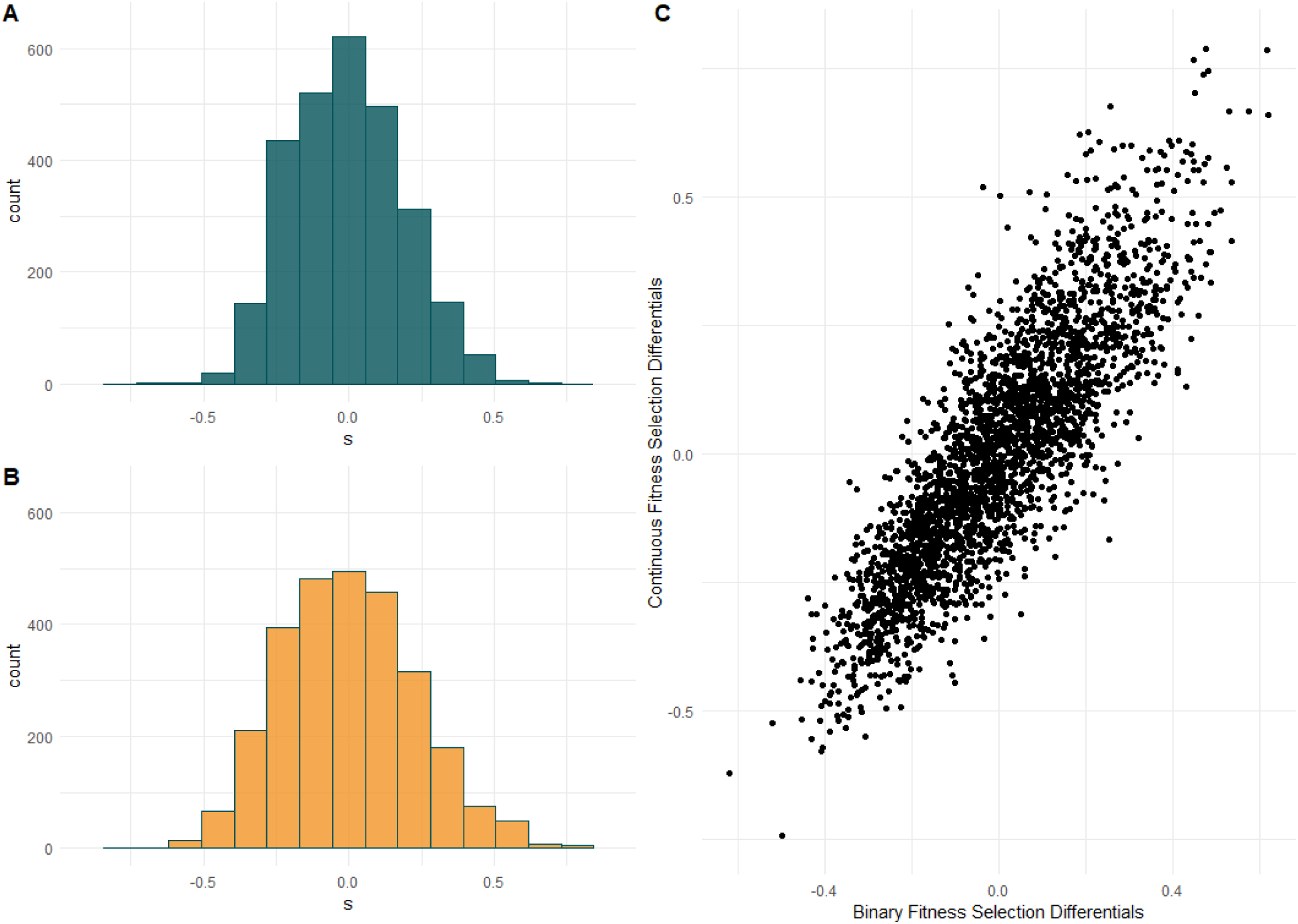
Selection differentials for gene expression calculated using the binary fitness (set seed or did not set seed) and using the relative fitness in number of set seed. A) Histogram of the binary fitness selection differentials for standardized gene expression counts of 2753 genes. B) Histogram of the selection differentials as the covariance between relative fitness and standardized gene expression counts. C) The relationship between the selection differentials calculated using the two methods for each gene. The binary fitness selection differentials have a slightly narrower distribution but are highly correlated with the continuous fitness selection differential (Pearson’s correlation coefficient = 0.85, p < 0.001).

### Unsupervised modeling

The Principal Component Analysis (PCA) transformed the 2753 standardized gene expression counts from 96 individuals into 96 linear combinations of the standardized gene expression measures. The first 10 principal components (PCs) describe 50.62% of the variation in gene expression, and 68 PCs are required to explain >90% of the variation (recall that PCs are in descending order of the variance in gene expression they explain). We then regressed relative fitness on each PC individually to get the univariate regression p-value which we used to evaluate the model fit with varying numbers of PCs as predictor variables. We assessed model fits across different p-value cut-offs.Our best model included a total of 61 PCs with individual p-values <0.60, which minimized the RMSE (root mean-squared error) and MAE (mean absolute error), while maximizing the R-squared value (Figure S1). We note that these 61 PCs are *not* the first 61, and that the overall model which minimizes the RMSE and MAE contains PC axes that are not, on their own, significant predictors of fitness (Figure S2). The *caret* package, which we used to perform our repeat cross-validation regressions, uses the correlation between the predicted and observed response values as the R-squared value (Kuhn 2008). The model R-squared was 0.549, with an RMSE of 1.719 and MAE of 1.381, and included 61 PCs (Table 1).

**Table 1:** Metrics of model fit for both supervised and unsupervised methods. Gradient boosting metrics are calculated from the testing data only. PCA regression metrics are from repeat five-fold cross-validation, and R-squared is calculated as the correlation between the observed and predicted data.

| Model | Balanced Accuracy | Sensitivity | RMSE | R-squared |
| --- | --- | --- | --- | --- |
| Gradient Boosting | 0.737 | 0.545 | - | - |
| PCA regression | - | - | 1.719 | 0.549 |

We then transformed the regression coefficients for the PCs used in the final model back to the standard-deviation scaled gene expression counts using the rotations from the PCA (Chong et al. 2018) to evaluate the strength and distribution of gene expression selection gradients (β). We also calculated the standard error of the selection gradient for each gene’s expression (following Chong et al. 2018), and then evaluated whether the absolute value of the selection gradient was greater than 1.96 × its standard error (SE), following the standard formula for a 95% confidence interval (Sokal and Rohlf 1995). Many (∼59%) of the absolute selection gradients were greater than 1.96 × SE, suggesting that most of our estimates are significantly greater than sampling error, within the multivariate space spanned by these 61 PC axes. The distribution of β*’*s was centered on zero, with an approximately normal distribution, which suggests that there is directional selection for both increased and decreased gene expression (Fig. 2). Overall, the magnitude of β*’*s was very small: an order of magnitude smaller than the significant β*’*s estimated for macroscopic traits in the same experiment, as reported by Henry and Stinchcombe (2023; Fig. 2B). The only β for macroscopic, traditional traits that was within the range of the gene expression selection gradients was for anther-stigma distance, which was not significant. Given that these selection gradients are on individual genes and not the combined outcome of many genes, as is the case for the quantitative traits studied by Henry and Stinchcombe (2023), we expected the directional selection gradients on each gene’s expression individually to be small. In addition, if any of the expressed genes are themselves causally associated with phenotypic traits under selection (i.e., expression → trait → fitness), we expect a small estimate for expression’s net effect on fitness (expression → fitness), due to it being the product of two small numbers.

**Figure 2:**
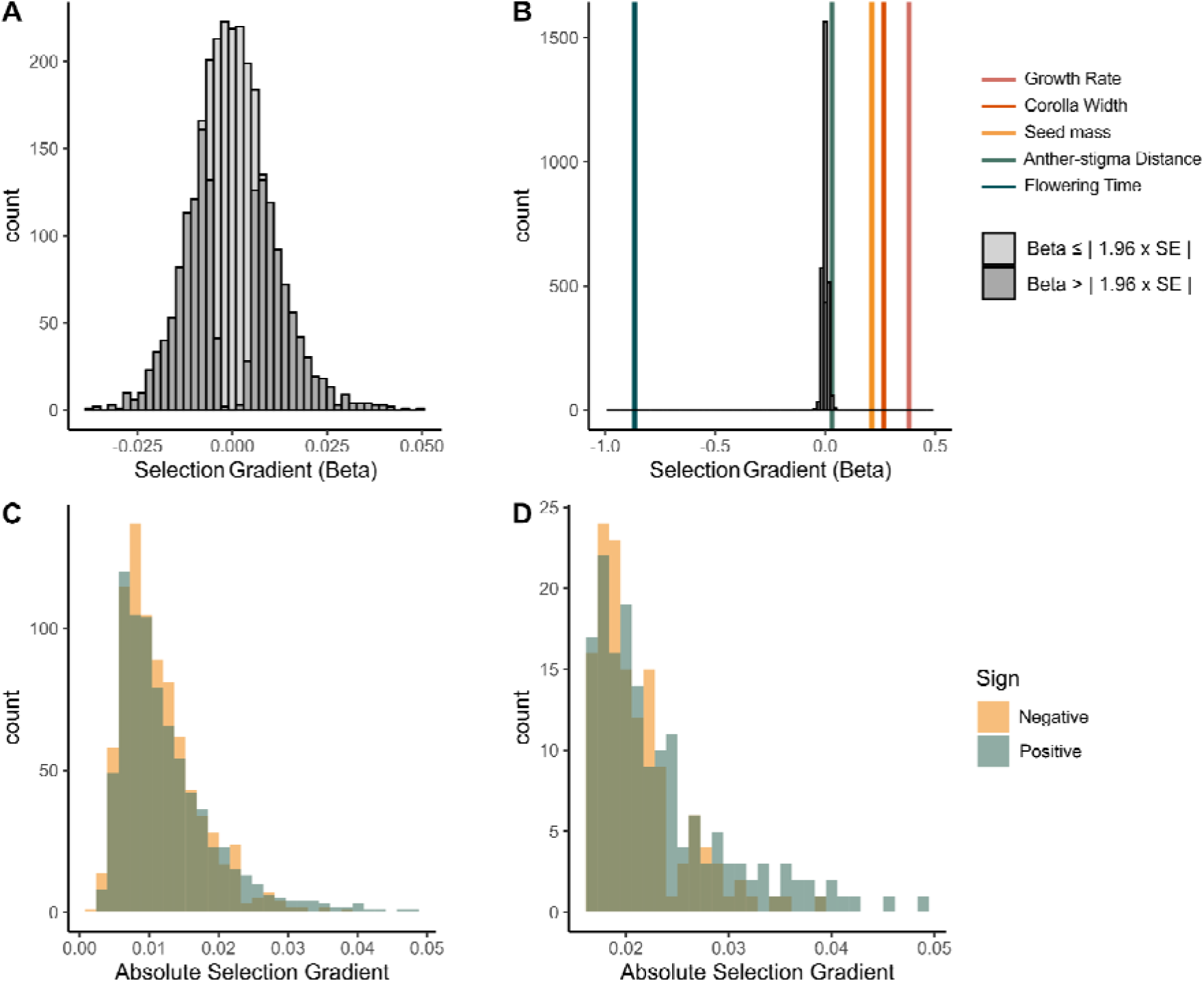
Distribution of selection gradients for gene expression of the 2753 genes. A) Selection gradients shaded by whether they were greater than 1.96 x their standard error. B) The same distribution of selection gradients plotted alongside the selection gradients of macroscopic phenotypes. C) Selection gradients plotted as their absolute values, yellow representing the negatively signed selection gradients, and blue representing the positively signed selection gradients. D) Absolute selection gradients that are greater than 1.96 x their standard error.

### Supervised modeling

We tested a variety of hyperparameters through a grid search with five-fold cross validation for our supervised model. Our supervised machine learning model was a gradient boosting classifier, which fits weakly predictive decision trees, progressively adjusting the parameter weights to improve classification performance of misclassified observations. We tuned the hyperparameters of the model using a grid search with five-fold cross validation of the data. The best fitting model used a binomial deviance loss function, and the decision split quality was determined by Friedman adjusted mean-squared error (Friedman 2001). The models were regularized by using stochastic gradient boosting, which randomly subsamples the test data each iteration to reduce overfitting in the final ensemble model. We used 1000 estimators, a constant learning rate of 1.0, and subsampled 40% of the data each iteration. The final model had an out-of-sample balanced accuracy score of 0.737, with sensitivity to the successful seed set class of 0.545 (Table 1, Figure S3). We extracted the 278 most important features based on their Gini importance from the model, with a cutoff of > 0.000677798, as the next 197 values were tied.

### GO Analysis

Once we obtained a subset of most important genes for each model, we performed BLAST searches on gene sets using an *Arabidopsis thaliana* genome (Lamesch et al. 2012) as a reference, and then mapped and annotated GO terms. We generated combined GO maps for our gene subsets, filtering out intermediates, for each model. We found 11 GO terms common across our two models, in addition to the higher level “biological process” term (Table 2). The shared terms are involved in developmental and metabolism, and various stimulus responses. These common terms highlight the importance in the response to both plant enemies (defense response, defense response to other organisms, and response to biotic stimulus), and the abiotic environment (response to light stimulus, and development terms). More broadly, non-identical but related GO terms common across the models were associated with defense and stress response (GO:0006950, GO:0009737, GO:0051707, GO:0009620), metabolism (GO:0051171, GO:0060255, GO:0016071, GO:0080090, GO:0034641, GO:0006518,GO:0008152, GO:0032787, GO:0044238, GO:0016070, GO:0071704, GO:0006629) and reproductive structure development (GO:0048608, GO:0099402, GO:0009653, GO:0022414, GO:0048731, GO:0009791) (see Supplemental Material, Table S2 – Table S3).

**Table 2:**
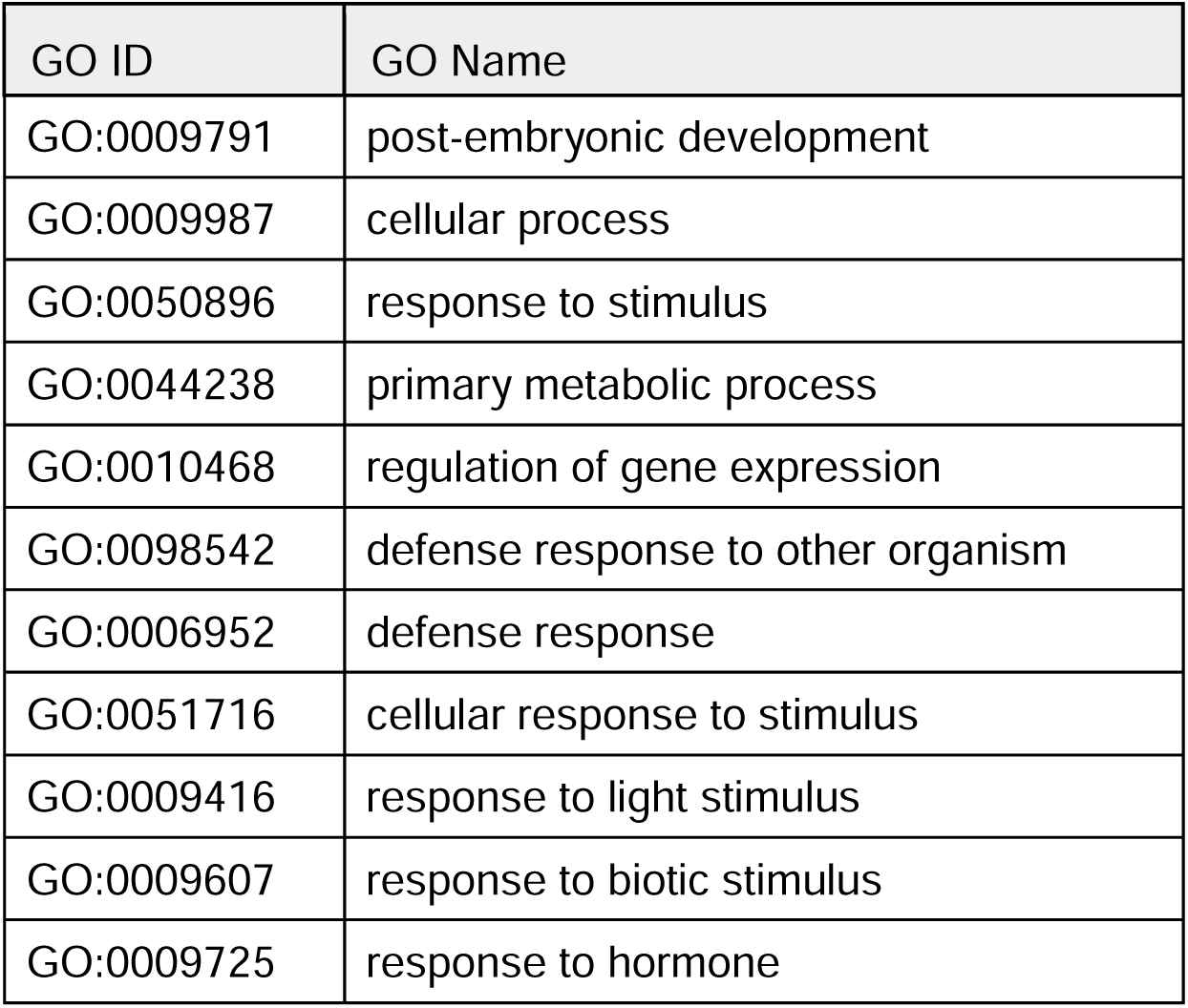
Shared GO terms from GO mapping of gene subsets. Gene subsets were composed of the most important genes in each mode, which were included in a BLAST search and annotated with GO terms. Response terms dominate, along with developmental and metabolic terms. The GO term for light response is also shared.

| GO ID | GO Name |
| --- | --- |
| GO:0009791 | post-embryonic development |
| GO:0009987 | cellular process |
| GO:0050896 | response to stimulus |
| GO:0044238 | primary metabolic process |
| GO:0010468 | regulation of gene expression |
| GO:0098542 | defense response to other organism |
| GO:0006952 | defense response |
| GO:0051716 | cellular response to stimulus |
| GO:0009416 | response to light stimulus |
| GO:0009607 | response to biotic stimulus |
| GO:0009725 | response to hormone |

We then used *goseq* to adjust for count bias in our gene set to determine which GO terms were significantly enriched in each model set, using *p* <0.10 as our threshold. Two GO terms were shared between our models, “seed maturation” (GO:0010431) and “vegetative to reproductive phase transition of meristem” (GO:0010228) The two sets also included similar terms related to photosynthesis, seed maturation and dormancy, stress, and heavy metals (see Supplemental Material, Table S4 – Table S5).

### Shared Important Genes

There were 29 shared genes in the sets of most important genes for each model. For the 29 genes that are among the top ranked in the PC regression and identified by the gradient boosting classifier, we observed that the values of selection differentials (median = 0.17, range = [-0.44, 0.79] ) and selection gradients (median = 0.01, range = -0.04, 0.03]) for these genes were bimodal, with fewer instances of values close to zero relative to the distribution of all of the genes that we studied (Fig 1, Fig 2, Fig. S5, Fig. S6). These data suggest that the gradient boosting classifier is identifying genes for further study and exploration that show the strongest patterns of phenotypic selection. Of particular note was that in this top 29 genes identified by each method, GO terms related to heavy metals were apparent in each. Consequently, we next turned to the heavy metal concentrations of the soils at our field site.

### Heavy Metal Analysis of Soils

None of the metals in the screen (Table S8) were above government agricultural guidelines (Ontario Ministry of the Environment 2011) and were largely below the average for agricultural fields in the area (Frank et al. 1976). While our sample of plant lines was from several states, heavy metal concentrations in the field were also below the state-wide average for North Carolina which is in the heart of *I. hederacea’s* native range (Smith et al. 2008). Collectively, these data suggest that neither an induced response to contaminated soil at our field site, nor long term adaptation to heavy metal soils in the native range, are contributors to the importance of the expression of LOC109174332 in predicting relative fitness. While heavy-metal associated plant proteins (HIPP) have clearly been found to be associated with heavy-metal tolerance and detoxification (hence their name), they have also demonstrated responses to drought and cold tolerance (Barth et al. 2009; de Abreu-Neto et al. 2013; Zschiesche et al. 2015; Zheng et al. 2023), which interact with plant growth and development (Zschiesche et al. 2015; Guo et al. 2021).

## Discussion

Connecting gene expression to fitness components can elucidate which genes are the targets of selection, but the high dimensionality, low sample size nature of gene expression data makes this goal computationally and statistically challenging. Machine learning (ML) techniques are well-suited to this sort of complex task, as feature selection, regularization to prevent overfitting (and thus poor prediction out of sample), and nonlinear functions are key aspects of many ML algorithms. In this study we successfully used gene expression in leaf tissue to predict the reproductive success of 96 *Ipomoea hederacea* individuals using statistical learning approaches. We find that after appropriate preprocessing of the data, a reasonable level of model accuracy can be attained using both supervised classification algorithms and semi-supervised regression, with our classification model predicting correctly over 70% of the test cases. Similarly, unsupervised dimensionality reduction followed by regression, had a correlation of 0.55 between the predicted and observed fitness values. Below, we discuss our results in light of the typical strength of selection gradients for gene expression, the insight of multiple analysis approaches, and GO categorization for understanding the relationship between expression and fitness components.

### Distribution and strength of selection on gene expression

We found appreciable evidence for phenotypic selection on gene expression, using variants of the same methods that have provided such a rich picture of selection on life history, phenology, morphology, and sexually selected characters (Hoekstra et al. 2001; Kingsolver et al. 2001; Kingsolver et al. 2012). There are, however, several caveats associated with our analyses. First, there is uncertainty in our estimates of selection. While we estimated standard errors, and 95% confidence intervals, for the selection gradients, the standard errors are only relevant in the portion of the data described by the PC axes included in our analyses; the formula for a confidence interval carries with it normality assumptions. Second, as with all selection analyses, there is the potential of missing traits not included in the model: it may be that expression of other genes (at other time points, or in other tissues) or other macroscopic traits are the true targets of selection, and that our selection estimates would change with their inclusion in our models. A knottier challenge is the possibility that both gene expression and relative fitness were responding to some unknown variable, and that the associations we detected were environmentally induced covariances (Rausher 1992; Stinchcombe et al. 2002). While we attempted to minimize this possibility using a spatially replicated, randomized design– and we detected little evidence of environmental covariances in estimates of selection on macroscopic traits (Henry and Stinchcombe 2023) – we cannot formally exclude the possibility. Finally, our estimate of fitness – lifetime seed production produced by an annual, largely selfing plant– omits other fitness components, such as germination success and over-winter seed survival in the seed bank. Many of these challenges are shared, by and large, by most estimates of phenotypic selection that have been gathered by evolutionary biologists. The problem of missing traits is inherent to the approach; many estimates of selection use fitness components or aspects of performance (e.g., size) rather than true fitness. Finally, most of the thousands of estimates of phenotypic selection in the literature (Hoekstra et al. 2001; Kingsolver et al. 2001; Kingsolver et al. 2011; Kingsolver et al. 20012; Siepielski et al. 2009; Siepelski et al. 2013) are also likely susceptible to environmental covariances. Replicated sampling of genotypes is one of the best approaches for addressing the environmental covariance problem, but in the context of gene expression, this is likely to be an expensive proposition.

With these caveats in mind, our results indicate that phenotypic selection on gene expression can be detected using moderate sample sizes, even in a non-model species. We used principal component analysis (an unsupervised machine learning approach) along with a repeat 5-fold cross validation approach to linear regression (a supervised machine learning approach) to estimate selection gradients. By back-transforming the selection gradients for the principal component scores to return the selection gradients for each individual gene’s expression, we can improve interpretability and further investigate the most relevant genes. We found that directional selection gradients for gene expression were overall symmetrically distributed around zero, and thus selection for both increasing and decreasing gene expression were approximately equal. The strength of selection was an order of magnitude lower than that of other traits measured in this field experiment (Henry and Stinchcombe 2023). Our finding was not unexpected, given low estimates of selection differentials in univariate analyses by Groen (2020) and by intuition — combined gene expression over time leads to higher level quantitative traits, and thus the individual expression components experience a fraction of the selection pressure compared to the total, end-point phenotype. We believe these results represent the first selection gradients for gene expression in the Lande-Arnold framework and illustrate the merits of PC regression for estimating selection on high-dimensional traits.

### Leveraging information from machine learning approaches

Both of our models had moderate predictive accuracy, which demonstrates the general applicability of machine learning in understanding selection on gene expression. The ensemble method of classification was able to predict with a moderately high degree of sensitivity given our limited data and had a balanced accuracy score of 0.737. Similarly, although not directly comparable, the regression model was able to explain a moderate amount of variation in relative fitness and had an r-squared value of 0.549. The noise inherent in tissue-level gene expression data paired with its high dimensionality and intrinsic modeling error can reduce confidence in the repeatability of any one analysis. However, comparing the results from disparate approaches, each optimized to reduce overfitting, improves the certainty in the overall importance of genes and biological processes which emerge repeatedly. For example, we found that genes identified as important for differentiating individuals that set seed or did not set seed through a gradient boosting classifier showed bimodal patterns of selection differentials and gradients, with either strong positive or negative coefficients. Beyond confirming that the expression of these genes is under strong phenotypic selection, our data suggest that a combined approach of using machine learning to identify important genes or features, in concert with estimating parameters for evolutionary prediction (selection differentials and gradients) may be a useful approach for understanding selection and evolution of gene expression.

### Implications from GO analysis

We found common GO terms through the union of the most influential genes in each of the models. These common terms suggest types of genes and gene networks whose expression is directly associated with fitness, and thus potentially experiencing selection. The terms which were common across all models from the GO mapping results were largely related to seed development, stress and defense responses, as well metabolic processes and light response. Growth rate is important for fitness, and nonlinear selection analysis has revealed that it has a slightly compensatory relationship with early flowering time (see Henry and Stinchcombe 2023), which is itself strongly selected for in *Ipomoea hederacea* grown in field environments (Simonsen and Stinchcombe 2010; Campitelli and Stinchcombe 2013b, Henry and Stinchcombe 2023). Metabolic and developmental processes are thus not unexpected terms to be relevant for predicting fitness related traits. Additionally stress and defense response genes were commonly important, suggesting that the individuals better able to respond to biotic and abiotic stressors such as water deficiency and fungal pathogens may be better able to successfully reproduce.

One of the genes important across all both models was a heavy-metal associated isoprenylated plant protein (LOC109174332) which, along with GO terms associated with abiotic stress lead us to test the site soil. However, the analysis of heavy metals in the soils of our field site suggest that, if anything, heavy metal concentrations are lower than regional averages or the typical soils from its native range, suggesting neither an induced response to metals nor long-term adaptation. GO enrichment analysis also suggests that genes related to iron ion starvation are more common than expected in the sets of influential genes from two of our models. Thus, it seems most likely that the significance of LOC109174332 lies in its role in mediating the stress responses and plant development in the field. These data suggest that expression responses to environmental challenges such as water deficiency and biotic interactions may be important for fitness in *Ipomoea hederacea* in the field.

Reproductive organ development GO terms were also found to be important in both models, which, given the observed strong selection on flowering time in this and other systems, might be expected. We found only two GO terms significantly enriched in our model subsets: seed maturation and vegetative to reproductive phase transition of meristem. The field site for this experiment is 400 km north of the observed northern range limit and the significance of these GO terms, along with shared non-identical GO terms associated with photosynthesis potentially highlights the importance of photosynthetic genes relevant for photoperiodic cues in development and flowering time (Shimizu et al. 2015). Our results here complement those of Groen et al. (2020), who also detected selection on photosynthesis in a field selection experiment on gene expression in rice under stressful environmental conditions.

### Future applications and conclusions

Here we have demonstrated the applicability of machine learning in detecting genes whose expression is associated with fitness in the field, despite a relatively small sample. The utilization of machine learning for associating genes with fitness related traits whilst including environmental influences has obvious applications for agricultural crop improvements. Finding genetic targets for increased seed or biomass production is a bedrock of modern plant breeding (Moose and Mumm 2008; International Wheat Genome Sequencing Consortium (IWGSC) 2018). Additionally, evaluating combinations of genetic features which have a concerted impact on fitness may be used for index selection, which may provide greater gains than improvement at single loci. More generally, similar applications of machine learning with gene expression could assist in identifying or validating associated higher-level phenotypes for trait-based studies or selection, especially for non-model organisms. Machine learning is a wide and developing field, with many potential applications in improving understanding of genetic and transcriptomic sequencing data in evolutionary biology. Despite noisy data, there is still enough signal of fitness differences that models were moderately successful with reasonable and affordable sample sizes, suggesting that measuring selection on gene expression with these and related tools is a promising approach. Overall, machine learning algorithms, and their combined results, are effective in understanding the selective importance of biological pathways and gene expression in a multivariate context, in understanding the strength and distribution of selection across the transcriptome and can be harnessed to find candidate genes and quantitative traits of interest for further study.

## Methods

### Study species and Natural History

We collected tissue from *Ipomoea hederacea* (Convolvulaceae), an annual vine commonly distributed throughout the eastern USA in agricultural fields, roadsides and other disturbed habitats. It is found as far north as New Jersey and southern Pennsylvania and ranges into Mexico. It exhibits latitudinal clines in both Mendelian and quantitative genetic traits (Bright and Rausher 2008; Simonsen and Stinchcombe 2010; Campitelli and Stinchcombe 2013a; Stock et al. 2014) and neutral genetic patterns consistent with metapopulation dynamics (Campitelli and Stinchcombe 2014). Germination typically occurs in early summer, and once plants begin to flower, they continue to do so until frost.

### Field Experimental Design and Sampling

For this study we subsampled tissue for gene expression from a larger field experiment (Henry and Stinchcombe 2023) where seeds were generated by self-fertilization in a common greenhouse environment. On 20 July 2021, we germinated scarified seeds into Pro-Mix BX mycorrhizae soil in 4” peat pots in a glasshouse at ambient temperature and light. Three days later, we transplanted pots into a recently ploughed old field at the Koffler Scientific Reserve (www.ksr.utoronto.ca), with pots placed into the ground flush with field soil; we soaked the individuals with water and provided no further or additional supplementation. We sampled tissue from 100 individuals from 56 populations (1-3 individuals per population, mean = 1.80), gathered from 10 states in *I. hederacea’s* eastern North American distribution, spanning ∼7° of latitude (33.017681° to 40.340767° N). The plants were left to grow naturally among emergent weeds and/or die naturally. A killing frost ended the experiment on 27 October 2021. For further details on the field experimental design and environment, see Henry and Stinchcombe (2023).

Based on preliminary results (*see Results*), we decided to sample soils at our field site for a panel of heavy metals. As a consequence of the time required for initial data analysis, soils were collected 1 year after the field experiment concluded, from the same field, which had been plowed on the same schedule as the year of our experiment. We collected soil from across five transects, homogenizing the soil sampled within each transect, and sent the samples to the Agricultural and Food Laboratories at the University of Guelph for evaluation. Levels of arsenic, cobalt, chromium, cadmium, copper, mercury, nickel, lead, and zinc were quantified.

### Tissue Collection and Extraction

We collected leaf tissue a single time, on 29 September 2021, 71 days after sowing. We chose this time point to coincide with the approximate midpoint of the life cycle when slightly more than half of plants were flowering. The spatial block from which we sampled leaf tissue started with 342 seeds planted; at the time of sampling 33 plants had died (9.6%), and 37 had never germinated (10.8%). From the remaining available, we chose individuals using stratified random sampling across the population of origin. We did not have *a priori* hypotheses about particular genes, or that expression in leaf tissue (versus roots, meristems, vascular tissue, etc) was specifically important in determining fitness, but rather sought to obtain a multivariate sample of the transcriptome at the approximate midpoint of the life cycle. In this way, our estimates of gene expression traits are analogous to other studies of phenotypic selection that measure a suite of correlated traits at one moment in time (e.g., anti-herbivory traits, size, branch architecture, and phenology). We sampled whole leaves approximately 2.5 cm in diameter and placed them into RNAse-free 2 mL microtubes with a 6mm sterile glass bead. We immediately submerged the samples in liquid nitrogen and transferred them into a cooler containing dry ice and excess liquid nitrogen until tissue collection was complete. We stored samples in a -80 °C freezer until we performed extractions in November 2021. We extracted mRNA using Qiagen Plant RNEasy Extraction kits and followed the standard protocol, using a TissueLyser to prepare samples. Genome Quebec performed NEB mRNA stranded library preparations and Illumina library quality control, prior to sequencing using paired-end Illumina mRNA sequencing (NovaSeq6000 S4 PE100 Sequencing). Of the 100 samples collected, 97 were successfully sequenced, representing 55 populations. Of those 97 individuals, 58 had flowered by leaf tissue collection, 86 had flowered by the end of the experiment and 26 had set seed by the end of the experiment, approximately 1 month later (27 October 2021). We examined the samples for quality and contamination using FastQC, summarized with MultiQC. The mean Phred score across all reads and samples was above 35, and no primer contamination was detected. The mean number of reads per sample was 31.7 million (range: 18-45.4 million).

### Data processing

To align the raw reads we performed a two-step STAR alignment (Dobin et al. 2013) using the *Ipomoea nil* genome (Hoshino et al. 2016); *I. nil* is the sister species to *I. hederacea* (Miller et al. 1999; One Thousand Plant Transcriptomes Initiative 2019), with variable inter-crossability between the two, mediated by style length (Esserman 2012). The genome indexes for *I. nil* were generated with STAR default parameters except for the ‘genomeSAindexNbases’ parameter, which was reduced from the default 14 to 13, given the size of the genome (Dobin et al. 2013). On average 92% of the reads were mapped successfully to the *I. nil* genome. We then used gffread (Pertea and Pertea 2020) to generate a transcript annotation file for use in Salmon (Patro et al. 2017), which we ran to quantify gene counts from the aligned samples. As we needed to compare across samples, we imported our quantified gene counts into R using *tximport* to compile the samples into a single data frame and transformed the read counts using the Trimmed-M Means transformation (Robinson and Oshlack 2010) using the *edgeR* package (Robinson et al. 2010; McCarthy et al. 2012; Chen et al. 2016). We then filtered out genes with very low expression where we would not expect to be able to detect differences between individuals that never flowered and those which flowered and successfully set seed; we did so using the *filterByExpr()* function in the edgeR package (Chen et al. 2016), which reduced the number of genes in our dataset to 2753. As we had two groups (set seed and did not set seed), and our set seed group had 26 individuals, we defined “very low expression” as genes where 10 counts per million were observed less in less than 19 samples (the function in edgeR finds this value as 70% of the smallest group size).

It is common to correct estimates of gene expression for batch effects or effects due to the time of day of collection (Leek et al. 2012; see, e.g., Josephs et al. 2020). We elected not to do so for several related reasons. First, plants were planted in the field in randomized order, and as such tissue collection was in random order with respect to population of origin, genotype, and other known and unknown features of the samples. Second, our goal is to evaluate the relationship between gene expression, which may include time-of-day effects on the order of minutes to hours, and relative fitness, estimated from the cumulative number of seeds set by the end of the experiment, four weeks later; fitness itself reflects ∼100 days of development, life history, and growth. Rather than focusing on individual genes or results from single models, we concentrate on broad patterns in the intersection of different types of models, each implemented with permutation testing and cross-validation. Finally, preliminary analysis suggested that only one of 96 principal components of gene expression showed a significant rank correlation with order of collection; this PC was not predictive of relative fitness, and was not included in our analyses (*see below*).

## Analytical Methods

### Unsupervised modeling

We first performed principal component regression for dimensionality reduction without filtering out features. We standardized the data such that the transformed gene expression counts such that each gene had a standard deviation = 1 and mean = 0. We removed one outlier, which had set 86 seeds (mean seed set = 1.75, range = [0 – 24], excluding the outlier), to improve prediction. We calculated relative fitness for the remaining 96 individuals, by dividing the individual seed set by the mean seed set. Individuals that set no seeds were retained, having relative fitness of zero (Figure S4).

We performed a principal component analysis on the transformed and standardized data using *prcomp* from the *stats* package in R (R Core Team 2022). To determine which principal components (PCs) to include in the model we took a supervised regression approach, as lower PCs may still possess variation which may predict relative fitness well (Chong et al. 2018). We eliminated the last PC as the variance it described was approaching zero and was likely to introduce multicollinearity (Jolliffe 2002).

We first ran simple linear regression with relative fitness regressed on the PC scores of each individual and compiled the p-values of each principal component regression. We then sorted the PCs based on those p-values and used a 100-repeat 5-fold cross-validation regression with PCs of increasing p-values to determine the optimal number of PCs to include. We used the *train* function in the *caret* package to do so, saving the model parameters and performance measure of each iteration. We evaluated the error rates (both root mean squared and absolute error) and R-squared values of the models to determine the best model. Our model fitting procedure retained PCs based on their ability to predict relative fitness, and as such, PCs that explained relatively little variation in gene expression (the trailing principal components) but predicted fitness well were retained. We continued to add PCs until the R-squared, mean absolute error (MAE) and root mean squared error (RMSE) stopped improving.

To evaluate the importance of each gene we back-transformed the regression coefficients using the PCA rotations which resulted in the selection gradients (Chong et al. 2018), which account for selection on the expression of other genes. We then sorted the genes by their absolute coefficients to determine the 300 genes which demonstrated the strongest selection gradients, and thus were most influential in predicting relative fitness.

### Supervised modeling

We performed supervised analyses in Python using scikit-learn (Pedregosa et al. 2011). We removed the outlier and split the data into a training set (60%, or 57 samples) and test set (40%, or 39 samples). We again standardized the data to remove differences due to variance in expression among genes, we mean-centered and scaled the training data and then transformed the test data using the mean and standard deviation of the training data. For our supervised analyses we used classification algorithms, and as such classed our samples as having set seed (and thus having fitness > 0) or not (and having a fitness = 0).

We used a grid search to tune the hyperparameters (such as learning rate and number of iterations) of our models, which runs models with each combination of selected model hyperparameter values (the “grid”) and compares the fit of each model. We started with coarser grids (a wider range of sparse hyperparameter options) to search a larger parameter space and tuning further with finer-scale grids. To manage overfitting in the model we used five-fold cross-validation during the grid search on the training data. Model score was calculated using the balanced accuracy score, as we had far fewer samples which set seed and balanced accuracy accounts for uneven class sizes. The balanced accuracy score is the average accuracy score of each group, that is, the fraction of true positive results out of the total positive cases and the fraction of the true negative results out of the total negative cases. In our case, the “positive” case is the “set seed” class, and the “negative” case is the “no seed set” class. We then tested the model on the 40% of the data withheld for testing and examined out-of-sample balanced accuracy and sensitivity of the less-common class (individuals that successfully set seed). Sensitivity is calculated as the true positive predicted results divided by the total positive cases.

For our second classifier we used an ensemble method, which trains a large number of “weak learners” or models that are only marginally better than random, taken together to produce a “committee” model. We used a gradient tree boosting classifier, which is an ensemble method that uses some number of decision trees as weak learners, progressively adjusting weights to increase the importance of poorly classified observations from the previous tree, to build a final model weighted by the accuracy of each base model (Hastie et al. 2009). From this model we were able to directly extract the importance of genes, which is given as the Gini importance of the features in the model. We selected the top 278 features, as there were 197 ties in the importance of genes below this threshold.

### GO analysis

We then sought to characterize the types of genes which were important in predicting “success” (i.e., having set seed) in each of the models individually and which types were common among all models, using Gene Ontology (GO) biological terms. We also performed GO enrichment analysis on the subsets of genes from our analyses. To do so we used Blast2Go (Conesa et al. 2005), first to perform BLAST using blastx-fast of the genes to an *Arabidopsis thaliana* genome (Lamesch et al. 2012), then EMBL-EBI Interpro scan (Cantelli et al. 2022) using default options, followed by mapping the associated biological GO terms and annotation by Blast2Go. We subset the most important genes in each of the models, as described above and for each subset we repeated the BLAST, mapping and annotation of GO terms. We constructed GO term mapping for each set and extracted the union to evaluate common GO terms. We then used *goseq* (Young et al. 2010) correcting for read count, comparing the entire set of genes from our samples to use as a reference for GO enrichment analyses of each of the model sets, using Fisher’s Exact Test with a p-value of 0.05 as a filter to generate a set of over and under expressed GO terms. Again, we took the union of the results to look at commonly enriched GO terms.

## Author contributions

G.A.H. and J.R.S contributed to the design, analysis and writing of the manuscript. G.A.H. performed the data collection and implemented analyses.

## Data Availability

The data underlying this article available through the NCBI Sequence Read Archive https://www.ncbi.nlm.nih.gov/bioproject/PRJNA1030903/.

## Acknowledgements

We gratefully acknowledge financial support from NSERC Canada Discovery Grants (JRS, GAH), Weis-Zimmerman Graduate Fellowships in field biology (GAH), and the Swedish Collegium for Advanced Study (JRS). Comments from Stephen Wright, Jacqueline Sztepanacz, Art Weis, Megan Bontrager, and Joel McGlothlin improved the manuscript. We thank 3 anonymous reviewers and Kirk Lohmueller for helpful comments. Computational support was provided by the Digital Research Alliance of Canada. We thank Kate Brown and Radana Molnarova for support in the field at the Koffler Scientific Reserve.

