## Supplementary materials for "Predicting fitness related traits using gene expression and machine learning"

**Supplementary material**
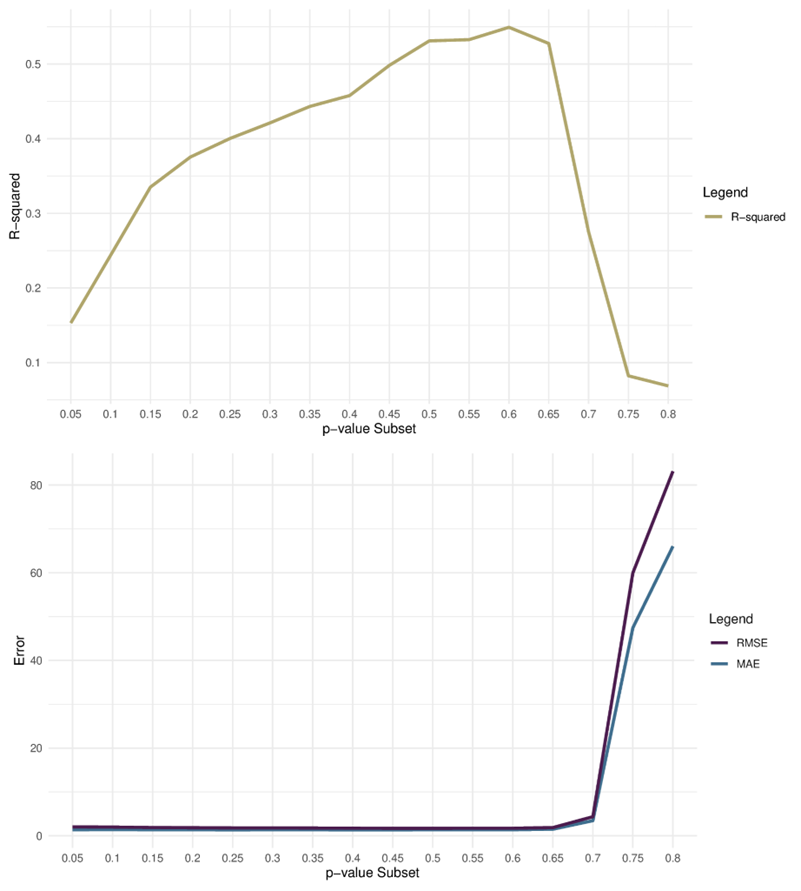

Figure S1: Principal component regression metrics. Relative fitness was regressed on each principal component individually, and then PCs were binned by p-value. Relative fitness was then regressed on subsets of the PCs based on the binned p-values. We chose p < 0.60 as the cut-off for PCs to include in the final model based on the balance of the R-squared value and error rates.

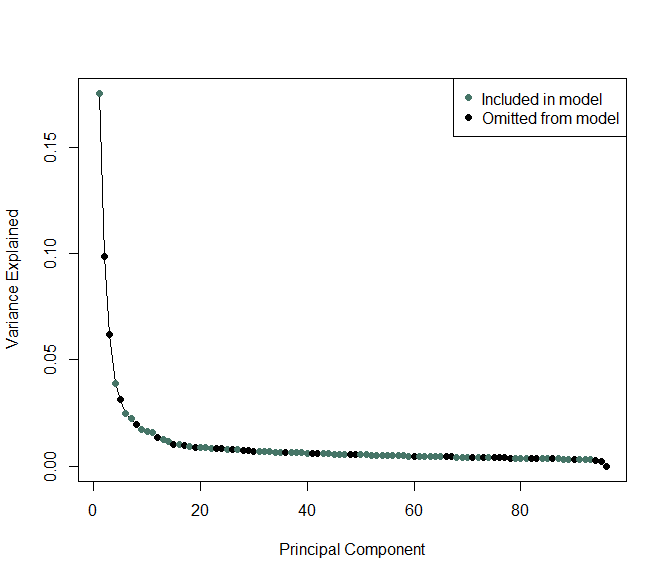

Figure S2: Scree plot of variance explained by each principal component of the full PCA on 2753 gene counts in 97 individuals. The PC’s coloured blue are those which we kept in our Principal Component Regression model, chosen via the p-values from individual regressions of fitness on each PC. We chose the p-value threshold that minimized root mean squared error and mean absolute error while maximizing r-squared, which in our case was p <= 0.60.

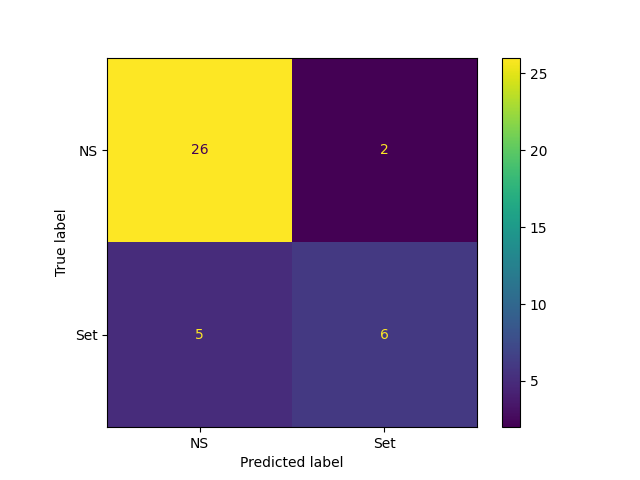

Figure S3: Gradient Boosting Classifier confusion matrix. NS label represents individuals that did not set seed, Set label represents individuals that did set seed. Predicted vs. observed labels for the testing data only. Precision, balanced accuracy, and sensitivity are calculated from these values.

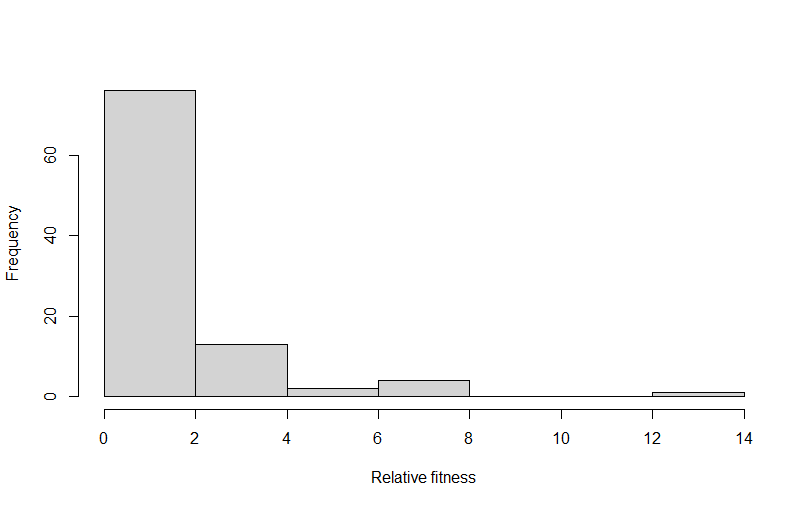

Figure S4: Histogram of the distribution of relative fitness in the 96 assayed individuals. Relative fitness was calculated as the individuals’s number of seeds produced scaled by the mean number of seeds ($x_{i} / \underline{x} )$.

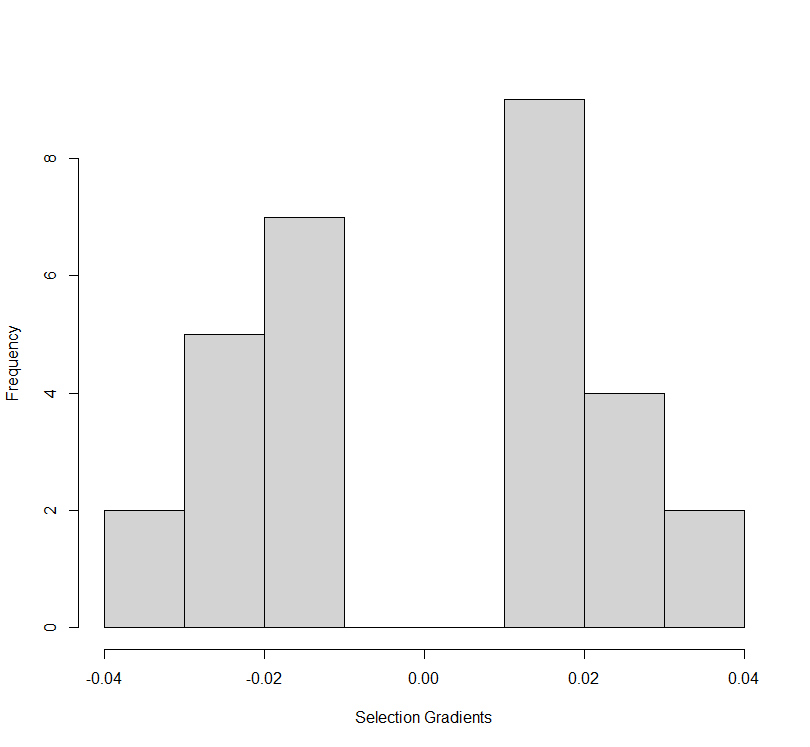
Figure S5: Selection gradients for the 29 shared most important genes in the PC Regression and Gradient Boosted Classifier models.

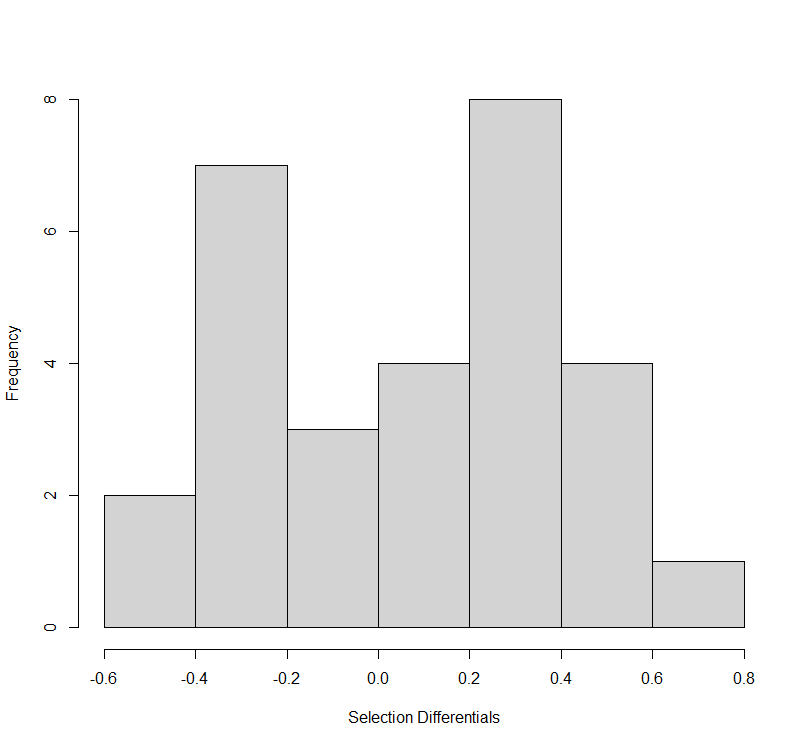

Figure S6: Selection differentials from continuous relative fitness (from seed number) for the 29 shared most important genes in the PC Regression and Gradient Boosted Classifier models.

Table S1: Model hyperparameters for gradient boosting classification model.

|  | Gradient Boosting Classifier | |
| --- | --- | --- |
|  | Parameter | Value |
|  | ccp_alpha | 0 |
|  | criterion | friedman_mse |
|  | init | None |
|  | learning_rate | 1 |
|  | loss | log_loss |
|  | max_depth | 10 |
|  | max_features | None |
|  | max_leaf_nodes | None |
|  | min_impurity_decrease | 0 |
|  | min_samples_leaf | 1 |
|  | min_samples_split | 2 |
|  | min_weight_fraction_leaf | 0 |
|  | n_estimators | 1000 |
|  | n_iter_no_change | None |
|  | random_state | None |
|  | subsample | 0.4 |
|  | tol | 0.0001 |
|  | validation_fraction | 0.1 |
|  | verbose | 0 |
|  | warm_start | FALSE |

Table S2: Filtered GO terms from combined GO mapping of Gradient Boosting Classifier gene subset. GO terms are nested such that terms with lower level numbers are subdivided into associated terms with higher level numbers. Node score describes the annotation density of the GO term, based on distance to connected terms and number of sequences. Table continues on next page and is available as an Excel spreadsheet.

| Level | GO ID | GO Name | Node score | #Seqs |
| --- | --- | --- | --- | --- |
| 1 | GO:0008150 | biological process | 1110.68 | 221 |
| 2 | GO:0099402 | plant organ development | 60.59 | 29 |
| 2 | GO:0022414 | reproductive process | 68.15 | 43 |
| 2 | GO:0048731 | system development | 104.38 | 46 |
| 2 | GO:0009653 | anatomical structure morphogenesis | 38.73 | 22 |
| 2 | GO:0009987 | cellular process | 491.62 | 191 |
| 2 | GO:0050896 | response to stimulus | 409.5 | 114 |
| 2 | GO:0009791 | post-embryonic development | 70.51 | 36 |
| 2 | GO:0008152 | metabolic process | 563.18 | 150 |
| 2 | GO:0065007 | biological regulation | 182.56 | 85 |
| 2 | GO:0051707 | response to other organism | 104.31 | 42 |
| 2 | GO:0050789 | regulation of biological process | 152.74 | 77 |
| 2 | GO:0006810 | transport | 66.02 | 36 |
| 3 | GO:0009416 | response to light stimulus | 32.3 | 23 |
| 3 | GO:0032787 | monocarboxylic acid metabolic process | 37.59 | 22 |
| 3 | GO:0009620 | response to fungus | 33.2 | 23 |
| 3 | GO:0051716 | cellular response to stimulus | 80.05 | 44 |
| 3 | GO:0098542 | defense response to other organism | 58.71 | 32 |
| 3 | GO:0019438 | aromatic compound biosynthetic process | 47.28 | 37 |
| 3 | GO:0044238 | primary metabolic process | 214.1 | 118 |
| 3 | GO:0009607 | response to biotic stimulus | 107.31 | 43 |
| 3 | GO:0016070 | RNA metabolic process | 64.98 | 40 |
| 3 | GO:0010468 | regulation of gene expression | 59.02 | 31 |
| 3 | GO:0009058 | biosynthetic process | 177.01 | 82 |
| 3 | GO:0071840 | cellular component organization or biogenesis | 91.55 | 58 |
| 3 | GO:0071704 | organic substance metabolic process | 339.38 | 132 |
| 3 | GO:0010033 | response to organic substance | 86.06 | 41 |
| 3 | GO:0006952 | defense response | 77.72 | 38 |
| 3 | GO:0009725 | response to hormone | 63.37 | 31 |
| 4 | GO:0009059 | macromolecule biosynthetic process | 51.29 | 37 |
| 4 | GO:0044085 | cellular component biogenesis | 32.32 | 24 |
| 4 | GO:0006629 | lipid metabolic process | 31.83 | 22 |
| 4 | GO:1901362 | organic cyclic compound biosynthetic process | 53.3 | 41 |

Table S3: Filtered GO terms from combined GO mapping of Principal Component Regression gene subset. GO terms are nested such that terms with lower level numbers are subdivided into associated terms with higher level numbers. Node score describes the annotation density of the GO term, based on distance to connected terms and number of sequences. Table continues on next page and is available as an Excel spreadsheet.

| Level | GO ID | GO Name | Node score | #Seqs |
| --- | --- | --- | --- | --- |
| 1 | GO:0008150 | biological process | 835.26 | 224 |
| 2 | GO:0034641 | cellular nitrogen compound metabolic process | 130.55 | 98 |
| 2 | GO:0006518 | peptide metabolic process | 40.89 | 26 |
| 2 | GO:1901566 | organonitrogen compound biosynthetic process | 56.02 | 35 |
| 2 | GO:0019222 | regulation of metabolic process | 196.04 | 57 |
| 2 | GO:0006355 | regulation of DNA-templated transcription | 43.52 | 26 |
| 2 | GO:0051252 | regulation of RNA metabolic process | 44.98 | 34 |
| 2 | GO:0010467 | gene expression | 197.51 | 80 |
| 2 | GO:0006412 | translation | 38.02 | 23 |
| 2 | GO:0048608 | reproductive structure development | 63.66 | 38 |
| 2 | GO:0009791 | post-embryonic development | 93.21 | 50 |
| 2 | GO:0044238 | primary metabolic process | 272.06 | 144 |
| 2 | GO:0050896 | response to stimulus | 439.51 | 119 |
| 2 | GO:0050794 | regulation of cellular process | 145.92 | 77 |
| 2 | GO:0009987 | cellular process | 476.79 | 202 |
| 2 | GO:0006396 | RNA processing | 60.6 | 30 |
| 2 | GO:0051171 | regulation of nitrogen compound metabolic process | 67.74 | 46 |
| 2 | GO:0007165 | signal transduction | 53.67 | 31 |
| 2 | GO:0060255 | regulation of macromolecule metabolic process | 130.25 | 52 |
| 2 | GO:0098542 | defense response to other organism | 50.47 | 25 |
| 2 | GO:0032502 | developmental process | 181.14 | 78 |
| 2 | GO:0016071 | mRNA metabolic process | 38.5 | 22 |
| 3 | GO:0006950 | response to stress | 174.02 | 74 |
| 3 | GO:0009607 | response to biotic stimulus | 53.6 | 34 |
| 3 | GO:0010468 | regulation of gene expression | 89.33 | 41 |
| 3 | GO:0051716 | cellular response to stimulus | 87.9 | 47 |
| 3 | GO:0009725 | response to hormone | 93.85 | 45 |
| 3 | GO:0009628 | response to abiotic stimulus | 122.28 | 61 |
| 3 | GO:0042221 | response to chemical | 132.83 | 77 |
| 3 | GO:0080090 | regulation of primary metabolic process | 69.08 | 47 |
| 4 | GO:0009416 | response to light stimulus | 42.87 | 26 |
| 4 | GO:0006952 | defense response | 70.35 | 33 |
| 4 | GO:0009737 | response to abscisic acid | 35.22 | 22 |

Table S4: Goseq count corrected gene ontology terms for the top 278 most important genes in the Gradient Boosting Classifier model. P-value cutoff from Fisher’s Exact Test is 0.10. This table is also available as an Excel spreadsheet.

| Category | Over-represented pvalue | Under-represented pvalue |
| --- | --- | --- |
| carotene catabolic process | 0.00 | 1.00 |
| response to endoplasmic reticulum stress | 0.00 | 1.00 |
| xanthophyll catabolic process | 0.01 | 1.00 |
| abscisic acid-activated signaling pathway involved in stomatal movement | 0.01 | 1.00 |
| N-terminal protein amino acid modification | 0.01 | 1.00 |
| meiotic cytokinesis | 0.01 | 1.00 |
| negative regulation of developmental vegetative growth | 0.01 | 1.00 |
| negative regulation of leaf development | 0.01 | 1.00 |
| negative regulation of trichome patterning | 0.01 | 1.00 |
| regulation of seed dormancy process | 0.01 | 1.00 |
| response to iron ion starvation | 0.01 | 1.00 |
| positive regulation of gene expression | 0.01 | 1.00 |
| gibberellin biosynthetic process | 0.01 | 1.00 |
| regulation of cell population proliferation | 0.02 | 1.00 |
| fructose 1,6-bisphosphate metabolic process | 0.03 | 1.00 |
| regulatory ncRNA-mediated post-transcriptional gene silencing | 0.03 | 1.00 |
| negative regulation of gibberellic acid mediated signaling pathway | 0.03 | 1.00 |
| chlorophyll biosynthetic process | 0.03 | 0.99 |
| vegetative to reproductive phase transition of meristem | 0.03 | 0.99 |
| clathrin-dependent endocytosis | 0.03 | 1.00 |
| regulation of protein catabolic process | 0.03 | 1.00 |
| protein secretion | 0.03 | 1.00 |
| histone methylation | 0.03 | 1.00 |
| selenium compound metabolic process | 0.03 | 1.00 |
| positive regulation of programmed cell death | 0.03 | 1.00 |
| DNA repair | 0.04 | 0.99 |
| L-ascorbic acid biosynthetic process | 0.04 | 1.00 |
| hyperosmotic salinity response | 0.04 | 0.99 |
| seed maturation | 0.04 | 0.99 |
| protein quality control for misfolded or incompletely synthesized proteins | 0.05 | 1.00 |
| response to molecule of fungal origin | 0.06 | 1.00 |
| regulation of reactive oxygen species metabolic process | 0.06 | 1.00 |
| thiamine diphosphate biosynthetic process | 0.06 | 1.00 |
| negative regulation of seed germination | 0.06 | 0.99 |
| suberin biosynthetic process | 0.06 | 1.00 |
| thiamine biosynthetic process | 0.06 | 1.00 |
| response to copper ion | 0.07 | 0.99 |
| polar nucleus fusion | 0.07 | 0.99 |
| granum assembly | 0.08 | 1.00 |
| developmental process | 0.08 | 0.99 |
| gamete generation | 0.08 | 1.00 |
| cell death | 0.08 | 0.99 |
| organonitrogen compound metabolic process | 0.08 | 1.00 |
| pH reduction | 0.08 | 1.00 |
| mRNA transcription by RNA polymerase II | 0.08 | 1.00 |
| transcription by RNA polymerase II | 0.08 | 1.00 |
| SRP-dependent cotranslational protein targeting to membrane, signal sequence recognition | 0.08 | 1.00 |
| obsolete protein initiator methionine removal | 0.08 | 1.00 |
| glycoprotein catabolic process | 0.08 | 1.00 |
| protein deglycosylation | 0.08 | 1.00 |
| response to microbial phytotoxin | 0.08 | 1.00 |
| methionine biosynthetic process | 0.09 | 0.99 |
| intracellular potassium ion homeostasis | 0.09 | 0.99 |
| establishment of tissue polarity | 0.09 | 1.00 |
| photosystem II oxygen evolving complex assembly | 0.09 | 1.00 |
| 7-methylguanosine mRNA capping | 0.09 | 1.00 |
| vesicle fusion with Golgi apparatus | 0.09 | 1.00 |
| obsolete protein initiator methionine removal involved in protein maturation | 0.09 | 1.00 |
| meristem structural organization | 0.09 | 0.99 |
| regulation of gene expression by genomic imprinting | 0.09 | 1.00 |
| response to biotic stimulus | 0.09 | 1.00 |
| histone H2B ubiquitination | 0.09 | 1.00 |
| 'de novo' NAD biosynthetic process from aspartate | 0.09 | 1.00 |
| positive regulation of sulfur metabolic process | 0.09 | 1.00 |
| ribosomal small subunit biogenesis | 0.09 | 1.00 |
| regulation of monoatomic anion transport | 0.10 | 1.00 |
| L-proline biosynthetic process | 0.10 | 1.00 |
| photosynthetic electron transport in photosystem II | 0.10 | 1.00 |

Table S5: Goseq count corrected gene ontology terms for the top 300 most important genes in the Principal Component Regression model. P-value cutoff from Fisher’s Exact Test is 0.10. This table is also available as an Excel spreadsheet

| Category | Over-represented pvalue | Under-represented pvalue |
| --- | --- | --- |
| histone H3-K9 deacetylation | 0.00 | 1.00 |
| protein import into nucleus | 0.00 | 1.00 |
| vegetative to reproductive phase transition of meristem | 0.00 | 1.00 |
| nucleosome assembly | 0.01 | 1.00 |
| RNA splicing | 0.01 | 1.00 |
| mRNA processing | 0.01 | 1.00 |
| jasmonic acid and ethylene-dependent systemic resistance | 0.01 | 1.00 |
| regulation of catalytic activity | 0.01 | 1.00 |
| phosphorylation | 0.01 | 1.00 |
| regulation of mRNA splicing, via spliceosome | 0.01 | 1.00 |
| regulation of leaf senescence | 0.01 | 1.00 |
| negative regulation of cell population proliferation | 0.01 | 1.00 |
| regulation of DNA endoreduplication | 0.01 | 1.00 |
| regulation of timing of transition from vegetative to reproductive phase | 0.01 | 1.00 |
| negative regulation of DNA recombination | 0.01 | 1.00 |
| positive regulation of stem cell population maintenance | 0.02 | 1.00 |
| mRNA stabilization | 0.02 | 1.00 |
| N-glycan processing | 0.02 | 1.00 |
| ribosomal subunit export from nucleus | 0.02 | 1.00 |
| chromosome condensation | 0.02 | 1.00 |
| regulation of translation | 0.02 | 1.00 |
| primary miRNA processing | 0.02 | 1.00 |
| deadenylation-independent decapping of nuclear-transcribed mRNA | 0.02 | 1.00 |
| positive regulation of cytoplasmic mRNA processing body assembly | 0.02 | 1.00 |
| auxin biosynthetic process | 0.02 | 1.00 |
| signal peptide processing | 0.02 | 1.00 |
| DNA-mediated transformation | 0.02 | 1.00 |
| seed morphogenesis | 0.03 | 1.00 |
| plant ovule development | 0.03 | 1.00 |
| intracellular protein transport | 0.04 | 0.99 |
| DNA-templated transcription initiation | 0.04 | 1.00 |
| proteasome-mediated ubiquitin-dependent protein catabolic process | 0.04 | 0.99 |
| seed maturation | 0.04 | 0.99 |
| stress granule assembly | 0.04 | 1.00 |
| regulation of gene expression | 0.05 | 0.99 |
| tetrahydrofolate interconversion | 0.05 | 1.00 |
| negative regulation of DNA-templated transcription | 0.05 | 0.98 |
| phosphatidylinositol phosphate biosynthetic process | 0.05 | 1.00 |
| chloroplast fission | 0.05 | 1.00 |
| response to gibberellin | 0.06 | 0.99 |
| glucosinolate metabolic process | 0.07 | 0.99 |
| salicylic acid catabolic process | 0.07 | 0.99 |
| unsaturated fatty acid biosynthetic process | 0.07 | 0.99 |
| ethylene biosynthetic process | 0.08 | 0.99 |
| response to toxic substance | 0.08 | 0.99 |
| regulation of seed germination | 0.08 | 0.98 |
| regulation of cell division | 0.09 | 0.99 |
| glutamate catabolic process | 0.09 | 1.00 |
| meiotic cell cycle | 0.09 | 0.99 |
| cellular response to calcium ion | 0.09 | 1.00 |
| eukaryotic translation initiation factor 4F complex assembly | 0.09 | 1.00 |
| formation of translation preinitiation complex | 0.09 | 1.00 |
| obsolete endonucleolytic cleavage involved in rRNA processing | 0.09 | 1.00 |
| petal vascular tissue pattern formation | 0.09 | 1.00 |
| sepal vascular tissue pattern formation | 0.09 | 1.00 |
| fatty acid homeostasis | 0.09 | 1.00 |
| cell population proliferation | 0.09 | 1.00 |
| post-transcriptional gene silencing | 0.10 | 0.99 |
| negative regulation of organ growth | 0.10 | 1.00 |
| plant ovule morphogenesis | 0.10 | 1.00 |
| response to ethylene | 0.10 | 0.97 |

Table S6: Soil analysis results from environmental metals screen. Southern Ontario agricultural mean and Ontario government guidelines for reference. The mean of all transects and the individual values for each transect through the field are presented. All values are given in mg/kg. This table is also available as an Excel spreadsheet

| Metal | S. Ontario Mean | Gov. Guideline | KSR Mean | KSR1 | KSR2 | KSR3 | KSR4 | KSR5 |
| --- | --- | --- | --- | --- | --- | --- | --- | --- |
| Arsenic | 6.3 | 11 | 1.82 | 2.1 | 1.9 | 1.8 | 1.7 | 1.6 |
| Cadmium | 0.56 | 1 | 0.15 | 0.14 | 0.14 | 0.17 | 0.16 | 0.14 |
| Chromium | 14.3 | 67 | 13.6 | 15 | 13 | 15 | 12 | 13 |
| Cobalt | 4.6 | 19 | 4.28 | 4.7 | 4.4 | 4.7 | 3.8 | 3.8 |
| Copper | 25.4 | 62 | 8.1 | 13 | 7.6 | 7.5 | 6.3 | 6.1 |
| Lead | 14.1 | 45 | 7.14 | 7.2 | 7.2 | 7.4 | 7.2 | 6.7 |
| Mercury | 0.08 | 0.16 | 0.0342 | 0.041 | 0.035 | 0.034 | 0.033 | 0.028 |
| Nickel | 15.9 | 37 | 9.08 | 11 | 9.2 | 9.5 | 7.9 | 7.8 |
| Zinc | 53.5 | 290 | 33.2 | 35 | 32 | 34 | 33 | 32 |
